# Atg1 modulates mitochondrial dynamics to promote germline stem cell maintenance in *Drosophila*

**DOI:** 10.1101/2022.09.08.507105

**Authors:** Minal S. Ayachit, Bhupendra V. Shravage

## Abstract

Mitochondrial dynamics (fusion and fission) are necessary for stem cell maintenance and differentiation. However, the relationship between mitophagy, mitochondrial dynamics and stem cell exhaustion is not clearly understood. Here we report the multifaceted role of Atg1 in mitophagy, mitochondrial dynamics and stem cell maintenance in female germline stem cells (GSCs) in *Drosophila*. We found that depletion of *Atg1* in GSCs leads to impaired autophagy (mitophagy) as measured by reduced formation of autophagosomes, increased accumulation of p62/Ref (2)P and accumulation of damaged mitochondria. Disrupting Atg1 function led to mitochondrial fusion in developing cysts. The fusion was a result of an increase in Marf levels in both GSCs and cysts, and the fusion phenotype could be rescued by overexpression of *Drp1* or by depleting *Marf* via RNAi in Atg1-depleted cyst cells. Interestingly, double knockdown of both *Atg1:Marf* affected ovariole size and the number of vitellogenic oocytes. While *Atg1:Marf* knockdown led to decrease in germ cell number. Strikingly, *Atg1:Marf* double knockdown leads to a dramatic loss of GSCs, GCs and a total loss of vitellogenic stages, suggesting a block in oogenesis. Overall, our results demonstrate that Drp1, Marf and Atg1 function together to influence female GSC maintenance and their differentiation into cysts.

**Research Highlights:**

1. Atg1, in addition to its role in mitophagy, influences mitochondrial dynamics during oogenesis through modulation of Marf.
2. Atg1 and Marf promote Germline stem cell maintenance in Drosophila.

## Introduction

Female GSCs have the ability to self-renew and differentiate into the egg. This ability depends on their interaction with the niche cells in *Drosophila*. GSC maintenance affects reproductive fitness and is crucial for the survival of the species[1]. Autophagy is an important homeostasis mechanism operational within all cells, including GSCs. Autophagy maintains proteostasis by removing toxic aggregates and redox homeostasis by removing damaged mitochondria via mitophagy. However, the molecular role of autophagy in the maintenance of GSCs is not understood[2]. Autophagy is a highly regulated and complex process that is initiated by Ulk1/Atg1, a serine-threonine kinase that phosphorylates downstream proteins that catalyze the formation of the autophagosome. Autophagosome fuses with the lysosome to form autolysosome, wherein the hydrolytic enzymes degrade the cargo and the contents released in the cytoplasm for recycling[3].

In metazoans, mitochondria are inherited maternally and are important for generating energy in the form of ATP. Mitophagy is crucial for the maintenance of GSCs by sensing and removal of dysfunctional mitochondria. Mitophagy is influenced by mitochondrial biogenesis, dynamics (fission and fusion), translocation (distribution) and degradation[4]. Perturbation in any of these processes can lead to an imbalance in redox homeostasis and mitochondrial dysfunction. If mitochondrial damage is beyond repair, the fusion machinery is inactivated, and mitochondrial fragmentation is promoted to facilitate mitophagy. Mitochondrial dynamics is controlled by activities of Dynamin-related protein 1 (Drp1), Mitochondrial assembly regulation factor (Marf) (Mitofusin 2 Mfn2 in mammals) and Optic atrophy 1 (Opa1). These GTPases hydrolyse GTP and use the energy to execute changes in mitochondrial morphology. Mfn2 is required for the fusion of outer mitochondrial membranes of mitochondria. Opa1 is important for the fusion of the inner mitochondrial membrane [5]. Fission requires GTPase Dynamin-related protein 1 (Drp1), which forms oligomers surrounding the mitochondria and cleaves the mitochondria. In *Drosophila*, loss of *Marf* and *Drp1* can lead to progressive loss of the female GSCs. Loss of *Marf* results in a reduction of pMad signaling, which is a self-renewal factor for female GSCs[6,7]. Mitophagy has also been demonstrated to be crucial for the development of the ovariole and oocyte. Mitochondria exhibit maternal inheritance, and it is vital for proper embryogenesis. Mitochondria with mutated mtDNA are eliminated during oogenesis by mitophagy and requires Atg1 and BNIP3 [8]. Despite these interesting observations, the relation between mitophagy, mitochondrial dynamics and stem cell exhaustion is not clearly understood.

We used *Drosophila* female GSCs to address the question of how mitophagy, mitochondrial dynamics, and stem cell maintenance are related. Female *Drosophila* harbor a pair of ovaries with 16-20 individual units called ovarioles within each ovary. The GSCs are located within the anteriormost structure called the germarium in each ovariole. It is surrounded by the niche cells that secrete factors required for maintenance of the ‘stemness’ while the daughter cells that exit the niche differentiate into cystoblast (CB). A single CB undergoes four asymmetric cell divisions to generate interconnected 16-celled cysts. One of the 16-celled cysts is chosen as the future oocyte, which is then sustained by the remaining 15 nurse cells[1]. The *Drosophila* ovary presents a simple and genetically tractable model to study mitophagy, mitochondrial dynamics and their role in stem cell maintenance.

Here we demonstrate the importance of Atg1 in the maintenance of basal autophagy and its role in mitophagy using *Atg1RNAi* and loss-of-function clones of *Atg1*. *Atg1* knockdown cyst cells also show a strikingly different mitochondrial morphology which implies a relation between the mitophagy machinery and the mitochondrial dynamics. Using a combination of genetics and cell biology, we demonstrate that Drp1, Marf and Atg1 collaborate to regulate stem cell maintenance in *Drosophila*.

## Materials and Methods

### Fly maintenance

All the transgenic fly stocks were maintained at 25°C on standard cornmeal sucrose malt agar. The following flies were used; *w; +; nosGal4VP16* (BL4937, RRID:BDSC_4937), *y sc v; +; Atg1-RNAi* (BL 35177, RRID:BDSC_35177), *, ywhsFLP12; NGT; Dr/TM6B, w; +;Drp1 RNAi, w:+; Marf RNAi, w;+; FIAsH-Flag-Drp1-HA, w;+; Marf-GFP; w;+; UAS-Drp1, w;+; UAS-Marf, Atg1Δ3D, ywhsFLP12; +; FRT80BhismRFP, ywhsFLP12; +; FRT80BubiGFP, w; +; FIAsH-Flag-Drp1-HA and w; +; Marf-GFP* were combined with *ywhsFLP12; NGT; Dr/TM6B, nosP mCherry-Atg8a nos 3’UTR, nosP-Ref(2)P-GFP* and *mito-roGFP2-Grx1* [2,9] were combined with w;+; *nosGAL4::VP16*.

### Generation of Atg1 clones

*FRT80BAtg1Δ3D* mutants crossed to FRT80BmRFP or FRT80BGFP were collected and fed on yeast for 2 days. The clones were generated using the FLP-FRT technique described previously.

### Immunostaining

Immunostaining was performed according to the protocol described in [2]. Antibodies and dilutions used; anti-CathepsinL 1:400 (Abcam Cat# ab58991, RRID:AB_940826); anti-ATP5α 1:250 (ThermoFisher Scientific Cat# 43-9800, RRID:AB_2533548), anti GFP 1:400 (Novus Cat# NB600-308, RRID:AB_10003058), anti-HA 1:1000 (CST Cat# 3724, RRID:AB_1549585), anti-Vasa 1:50 (DSHB Cat# anti-vasa, RRID:AB_760351), anti-Hts 1:50 (DSHB Cat# 1B1, RRID:AB_528070), anti-Lamin C 1:50 (DSHB Cat# lc28.26, RRID:AB_528339) and anti-pMad 1:50 (Abcam Cat# ab52903, RRID:AB_882596) 2° antibodies used: Alexa fluor 488 goat anti-rabbit 1:250 (Thermo Fisher Scientific Cat# A-11034, RRID:AB_2576217), Alexa fluor 555 goat anti-rat 1:250 (Thermo Fisher Scientific Cat# A-21434, RRID:AB_ 2535855), Alexa fluor 647 goat anti-mouse 1:250 (Thermo Fisher Scientific Cat# A-21236, RRID:AB_2535805), Alexa fluor 647 goat anti-rabbit 1:250 (Thermo Fisher Scientific Cat# A-21245, RRID:AB_2535813).

### Redox chemical treatment

Ovaries of *mito-roGFP2-Orp1* and *mito-roGFP2-Grx1* flies were dissected in Grace’s medium and washed in 1xPBS for 2 minutes. The redox treatment procedure was followed according to [2]

### Imaging and Analysis

All imaging was performed on Leica SP8 Confocal microscope using 63x oil objective. Images acquired were 8 bit, 1024×1024 pixel resolution at 100Hz scanning. Frame accumulation was performed with 6 frames for mCherry-Atg8a and GFP-Ref(2)P. Mito-roGFP2-Grx1 imaging and image analyses for all markers were performed as described in [2]. For mCherry-Atg8a puncta measurements in Atg1Δ3D (Atg1-/-) clones, an ROI was drawn around the GSCs and Atg1-/- clones. GSCs were identified by their close association with the niche cells and size of their nuclei while clones were recognized by absence of GFP. mCherry-Atg8a and GFP-Ref(2)P punctae were counted manually in RNAi mediated knockdown studies and the area of germarium were measured using ImageJ. Analysis of mitochondrial size was done by the MitoAnalyzer plugin. For colocalization analysis, we analyzed overlap of mCherry-Atg8a and CathepsinL, and GFP-Ref(2)P with CathepsinL puncta. Mitophagy flux was measured as colocalization between mitochondria (ATP5α) and lysosomes (Cathepsin L). Microsoft Excel was used for statistical analysis. Student’s T-Test of two samples assuming unequal variance was performed for all comparisons. GraphPad Prism7 was used to plot the graphs.

### DHE staining protocol

DHE staining and live imaging were performed according to manufacturer’s instruction (Invitrogen, USA).

### RNA isolation

20 pairs of ovaries were used for RNA extraction using the Trizol method as per manufacturer’s instruction (Invitrogen, USA).

### cDNA synthesis

1μg of tissue specific (ovary) total RNA was used for First strand cDNA synthesis using PrimeScript™ 1st strand cDNA Synthesis Kit (TakaraBio, India, Cat# 6110A).

### qPCR

qPCR was performed on EcoMax (Bibby Scientific, UK). The mean of housekeeping gene Tubulin was used as an internal control for normalization. The expression data was analysed using 2-ΔΔCT described by[10]

## Results

### Atg1 is necessary for basal autophagy and mitophagy during oogenesis

Atg1 is necessary and sufficient to induce autophagy in several tissues in *Drosophila*[3]. However, if Atg1 is required for autophagy in female GSCs is unclear. To address this, we examined the formation of autophagosomes in GSC-specific knockdown (KD) of *Atg1* by expression of a double-stranded inverse-repeat (IR) construct designed to target *Atg1* (UAS-*Atg1RNAi*) during oogenesis using *nanos-Gal4VP16*. Atg8a, the *Drosophila* ortholog of mammalian LC3, exhibits cytoplasmic localization but becomes incorporated into autophagosome when autophagy is induced and is visualized as punctate spots under fluorescence microscopy. GSCs and germ cells (GCs, also known as cyst cells) expressing *Atg1RNAi* had significantly fewer mCherry-Atg8a puncta as compared to the control (**Figure 1A-D and Supplementary Figure 1A, C**). Significant reduction of autophagy flux in *Atg1RNAi* germarium, as well as GSCs, was observed as seen by reduced colocalization between mCherry-Atg8a and CathepsinL (autolysosomes) (**Figure1C-D, Supplementary Figure 1D**). Further, we utilized FLP-FRT to generate *Atg1Δ30* mutant cell clones, resulting in tissue composed of control (either wild type or heterozygous *Atg1Δ30* /wild type, expressing GFP) and homozygous *Atg1 Δ30/Atg1 Δ30* (termed *Atg1-/-* henceforth, lacking GFP) mutant cells. Control cells contained several mCherry-Atg8a puncta, whereas homozygous *Atg1-/-* mutant cells (lacking GFP) displayed diffuse localization of mCherry-Atg8a (**Supplementary Figure 1E-E”, F**). These data suggest that Atg1 is necessary for autophagosome formation in the GSCs (including GCs).

**Figure 1.**
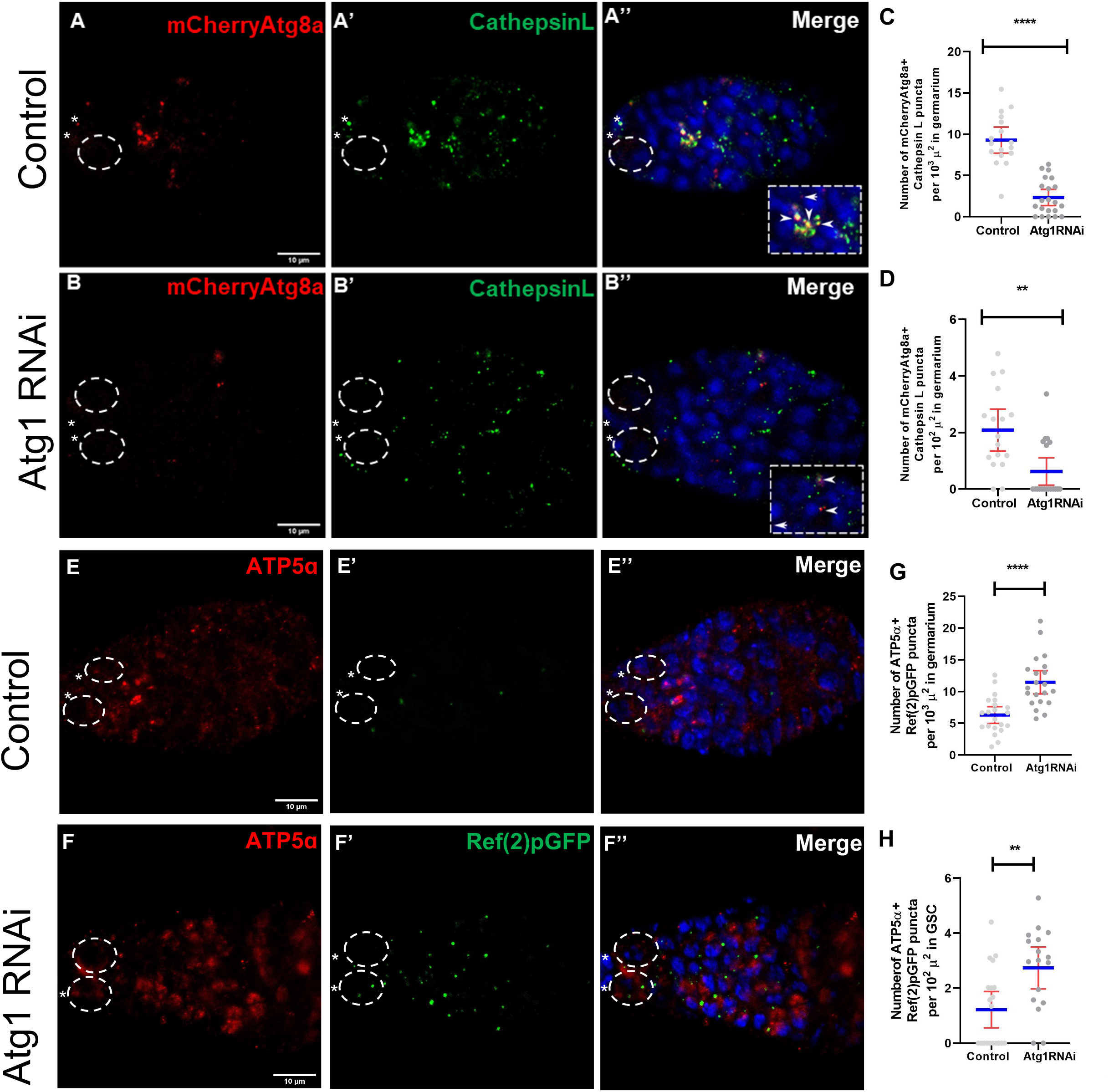
Atg1 is required for induction of autophagy during oogenesis. (A,B) Control (A-A”) and *Atg1RNAi* (B-B”) germarium showing mCherryAtg8a (A,B red) and CathepsinL (A’, B” green) staining. Inset in A”, B” shows enlarged region of colocalization (yellow puncta). Arrowheads point to colocalized punctae and arrows indicate mCherryAtg8a punctae (autophagosome). (C, D) Interleaved scatter graph representing overall decrease in autolysosomes (Cathepsin and mCherryAtg8a) positive structures in control and *Atg1RNAi* germarium (C) and GSCs (D). (E,F) Control (E-E”) and *Atg1RNAi* (F-F”) germarium ATP5α (E,F red) and Ref(2)pGFP (E’,F’ green) staining. Inset shows enlarged region of complete colocalization. Arrowheads point to colocalized punctae and arrows indicate Ref(2)PGFP punctae (cargo). (G,H) Interleaved scatter graph representing overall decrease in puncta/structures positive for ATP5α and Ref(2)PGFP in control and *Atg1RNAi* germarium (G) and GSCs (H). Nuclei is marked in blue (DAPI). Dotted ovals mark the GSCs and asterisk mark the cap cells. Scale bar-10μm. Error bars represent SD in red and the mean is represented in blue. n=20, **p < 0.05, **p < 0.01, ****p < 0.0001.

The *Drosophila* ortholog of p62 (SQSTM1), Ref(2)P binds ubiquitylated substrates, aids in their recruitment into autophagosomes, and is itself degraded in the autolysosomes. GSCs and cyst cells expressing *Atg1RNAi* accumulated GFP-Ref(2)P as compared to control germaria (**Figure 1E’-F’, Supplementary Figure 1B**). We also observed reduced colocalization between Ref(2)P and CathepsinL, suggesting a significant decrease in degradation of protein aggregates (**Supplementary Figure 1H-K**) This indicates that Atg1 is necessary for clearance of Ref(2)P during oogenesis via autophagy.

Autophagy is necessary for the removal of damaged mitochondria through mitophagy. p62/Ref(2)P is recruited to damaged mitochondria and has been shown to be necessary for mitophagy in mammals and *Drosophila* by facilitating the recruitment of damaged mitochondria to autophagosomes. Further, p62/Ref(2)P has been shown to be essential for PINK1-Parkin-mediated mitophagy, wherein decline of mitophagy was shown to be associated with increased co-localization with p62/Ref(2)P. As expected, we observed a significant number of punctate p62/Ref(2)P structures were found to colocalize with mitochondria in *Atg1RNAi* cells as compared to controls consistent with disruption of mitophagy (**Figure 1EH, Supplementary Figure 1G-G”**). Further, we measured mitophagy flux using colocalization of ATP5 alpha (red staining) and CathepsinL (green staining), followed by Pearson’s coefficient correlation. Mitophagy flux was found to be significantly reduced in *Atg1RNAi* expressing GSCs (and GCs) as compared to the controls, evident by reduced Pearson’s coefficient correlation (**Supplementary Figure 1L-O**). Taken together, these data suggest that mitophagy is disrupted in GSCs and cysts in *Atg1KD*.

Mitochondria within GSCs are elongated and become fragmented as cystoblasts differentiate into cysts cells[11]. In *Atg1KD*, mitophagy is disrupted in all cells within the germarium, but mitochondria were predominantly fused, particularly in the cyst cells, which was very surprising (**Figure 1E, F and Supplementary Figure 1L, M**). We did not observe hyperfused mitochondria in GSCs with *Atg1KD*. Next, we tested if this mitochondrial fusion phenotype was also observed in *Atg1-/-* mitotic clones. *Atg1-/-* mutant cyst cell clones (lacking GFP) possessed fused mitochondria in the germarium similar to *Atg1RNAi* (**Supplementary Figure 1G-G”**). Thus, disrupting Atg1 function leads to mitochondrial fusion during oogenesis, indicating that it controls mitochondrial dynamics in addition to mitophagy.

### Atg1 modulates Marf levels during oogenesis

Mitochondrial dynamics are regulated primarily by specific GTPases. Marf (Mitochondrial assembly regulatory factor) or Mitofusin 2, which regulates mitochondrial outer-membrane fusion, while *Opa1* mediates fusion of the inner membrane of mitochondria. Mitochondrial fission is catalyzed by *Drp1 (Dynamin-related protein 1*), which associates itself with the mitochondrial outer membrane and constricts mitochondria [6,7]. Mitochondrial fusion events can occur when Drp1 is reduced or when Marf and Opa1 levels increase. It is possible that cysts expressing *Atg1RNAi* have reduced activity of Drp1 or increased Marf activity. To test these possibilities, we monitored the levels of both Drp1 and Marf in *Atg1KD* cells. We utilized protein fusion reporters, Drp1-HA and Marf-GFP (Mfn2-GFP), that are expressed from their respective endogenous promoters[12]. No detectable difference in Drp1-HA levels was observed in *Atg1RNAi* expressing GSCs (and cyst cells) as compared to the controls (**Figure 2A-A”, B-B”, D, E and Supplementary data 2A-C**). We next tested if disrupting Drp1 function through the expression of *Drp1RNAi* could phenocopy *Atg1RNAi* mitochondrial fusion phenotype. Compared to the control, *Drp1RNAi* showed mitochondrial fusion in GSCs as well as in cyst cells (**Figure 2C-C”, F**). Strikingly, the mitochondrial phenotype observed in *Atg1KD* is similar to that of *Drp1KD*. Thus, the fusion of mitochondria in *Atg1KD* is consistent with the decrease in mitochondrial fission events caused by the depletion of fission regulator Drp1. Interestingly, our data also suggest that depleting *Drp1* leads to a reduction in *Atg1* transcription, indicating transcriptional regulation of *Atg1* by Drp1 (**Supplementary Figure 2D**).

**Figure 2.**
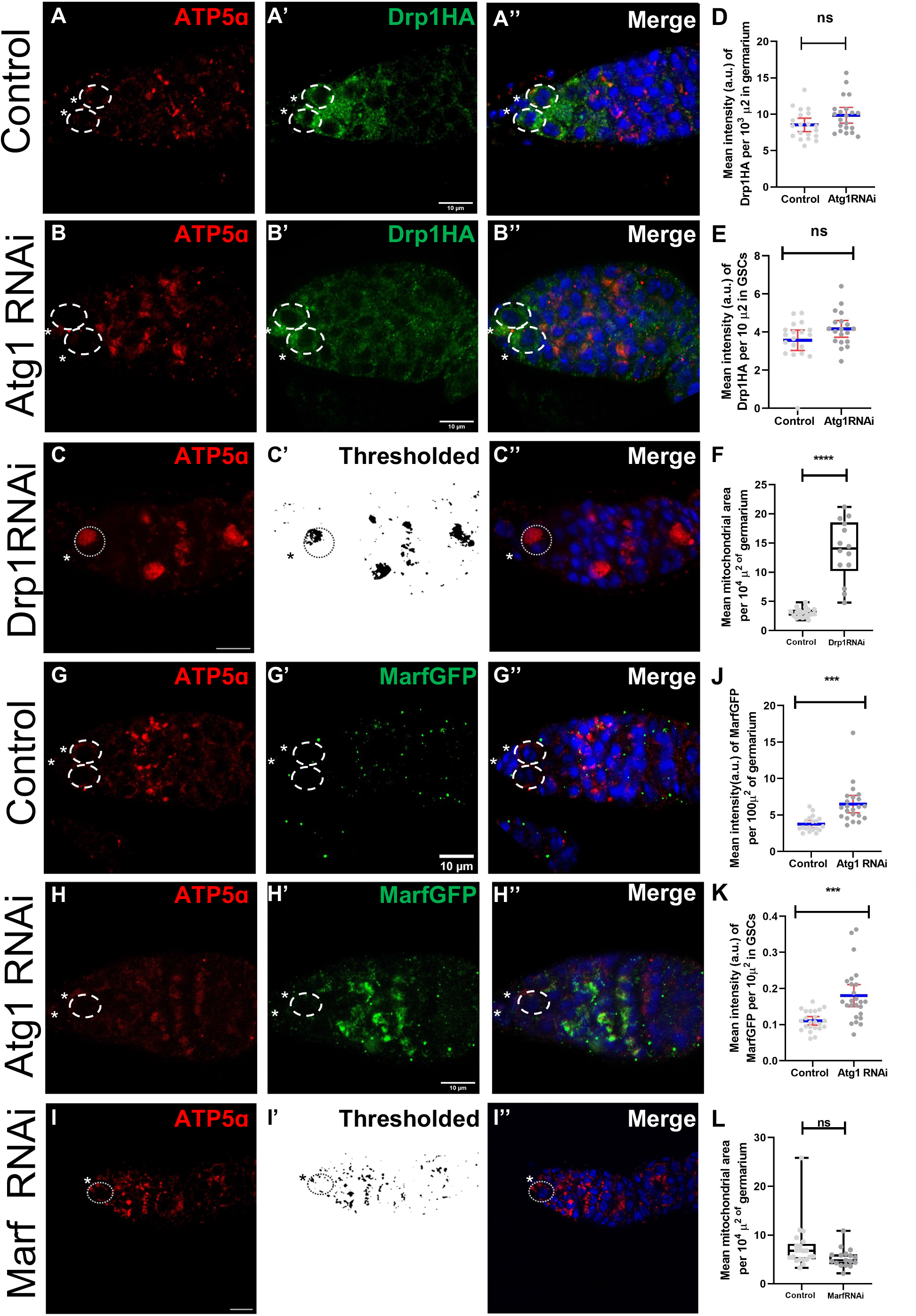
Atg1 regulates Drp1 and Marf and Drp1RNAi phenocopies Atg1RNAi mitochondrial phenotype. (A,B) Control (A-A”) and *Atg1RNAi* (B-B”) germaria immunostained with ATP5α (red) and HA (green) antibody to visualise HA tagged Drp1 protein.(C-C”) Germaria from *Drp1RNAi* stained with ATP5α (red) showing large clumped mitochondria (C), thresholded mitochondria (C’) and merged image with DAPI (C”). (D,E) Interleaved scatter graph showing the mean intensity of HA per unit area in germarium(C) and GSCs (D) of control and Atg1RNAi flies.(F) Box and whisker plot representing mean area of mitochondria measured per unit area of the germarium in C-C”.(G,H) Control (G-G”) and *Atg1RNAi* (H-H”) germarium immunostained for ATP5α (red) and GFP (green) and to visualise GFP tagged Marf protein.(J,K) Interleaved scatter graph showing the mean intensity of GFP per unit area in germarium (J) and GSCs (K) of control flies and *Atg1RNAi.(I-I”*) Germaria from *MarfRNAi* stained with ATP5α (red) showing fragmented mitochondria(I), thresholded mitochondria (I’) and merged image with DAPI (I”). (L) Box and whisker plot representing mean area of particles measured per unit area of the germarium in I-I”. Nuclei is marked in blue (DAPI). Dotted ovals mark the GSCs and asterisk mark the cap cells. Scale bar-10μm. Error bars represent SD in red and the mean is represented in blue. n=20, **p < 0.05, **p < 0.01, ****p < 0.0001.

Marf levels in *Atg1KD* GSCs and cysts were at least 2-fold higher as compared to the controls. These data are consistent with the fused mitochondrial phenotype observed in *Atg1RNAi* (**Figure 2G-G”, H-H”, J, K and Supplementary Figure 2D**). Further, we could detect a significant increase in Marf levels in region 1 and region 2 of *Atg1RNAi* expressing germaria as compared to the controls (**Supplementary Figure 2E**). Taken together, our results indicate that Atg1 is necessary to maintain fissed mitochondria in the cyst cells by regulating Marf levels in these cells. Further, we interfered with Marf function using RNAi-mediated KD and tested whether this affects mitochondrial morphology [6,7]. *MarfKD* caused fragmentation of mitochondria within germarium, including GSCs which is consistent with the reduction in fusion events upon depletion of fusion regulators (**Figure 2I, L and Supplementary Figure 2G**). Taken together, our data show that *Atg1KD* disrupts mitochondrial dynamics leading to fused mitochondrial morphology during early oogenesis.

### Atg1 genetically interacts with mitochondrial fusion machinery components

We next sought to test whether enhancing fission can rescue the *Atg1KD* mitochondrial fusion phenotype in cysts, given that Marf levels are elevated in *Atg1RNAi*-expressing germline cells. Fission could be enhanced by either elevating levels of Drp1 protein or by reducing Marf levels [6,7]. We hypothesized that by forcing mitochondrial fragmentation by overexpressing *Drp1* in *Atg1KD* will rescue the mitochondrial fusion phenotype. Similarly, interfering with mitochondrial fusion by reducing Marf (*MarfKD*) in combination with *Atg1KD* should lead to the rescue of the fused mitochondrial phenotype. We crossed *UAS-Drp1:Atg1RNAi* recombinants to *nanosGal4VP16* to co-express *Atg1RNAi* and *Drp1*, and assessed the mitochondrial fusion phenotype. *Drp1^OE^* in *Atg1KD* led to suppression of the mitochondrial fusion as observed by dispersed and punctate ATP5A1 staining in all GCs (**Figure 3A-C”, E and Supplementary Figure 3A, B, F**). The mitochondrial area was significantly reduced as compared to GCs expressing *Atg1RNAi* alone, suggesting that *Atg1* and *Drp1* genetically interact with each other to control mitochondrial dynamics. However, it is important to note that as compared to controls, the complete rescue of the mitochondrial phenotype was not observed.

**Figure 3.**
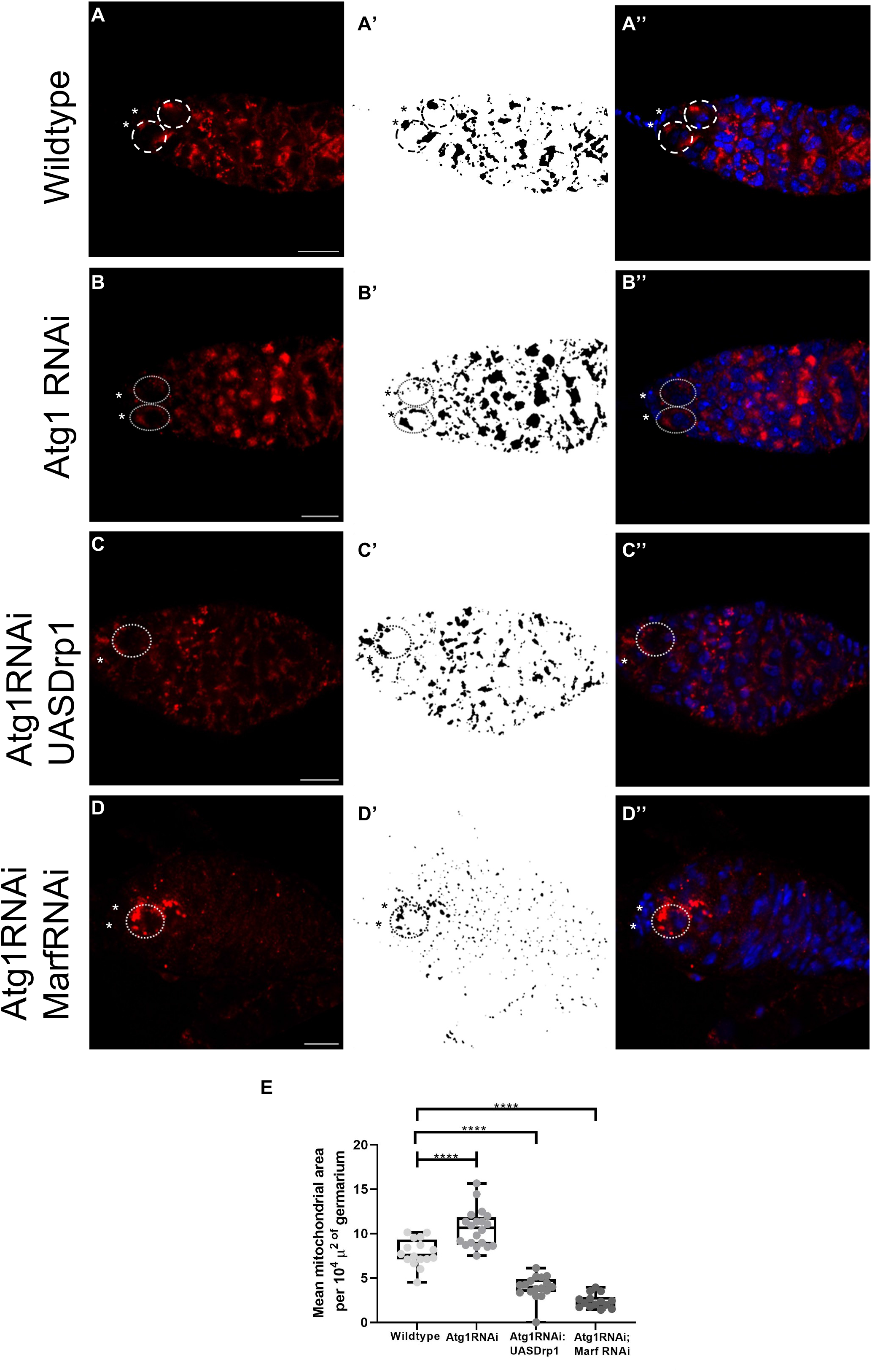
Atg1, Drp1 and Marf interact genetically to regulate mitochondrial dynamics during oogenesis. (A-D”) Germaria from wildtype, *Atg1RNAi, Atg1RNAi:UASDrp1* and *Atg1RNAi;MarfRNAi* stained with ATP5α (A-D red), thresholded mitochondria (A’-D’) and merged image with DAPI (A”-D”).(E) Box and whisker plot representing mean area of mitochondria measured per unit area of the germarium. Dotted ovals mark the GSCs and asterisk mark the cap cells. Scale bar-10μm. n=20, **p < 0.05, **p < 0.01, ****p < 0.0001.

We further tested if the mitochondrial fusion phenotype can be rescued by reducing Marf levels in *Atg1RNAi*. We co-expressed *Atg1RNAi* and *MarfRNAi* in all GCs. Previous studies have shown that a reduction in Marf levels within the GSCs leads to the fragmentation of mitochondria [6,7]. GSCs/GCs co-expressing *Atg1RNAi* and *MarfRNAi* exhibited fragmented mitochondria suggesting suppression of fused mitochondrial phenotype in GCs (**Figure 3A, B, D and E and Supplementary Figure 3A, C, F**). However, the total mitochondrial area was drastically reduced in combined KD of *Atg1:Marf*. Together these observations suggest that Atg1, Drp1 and Marf interact genetically to control mitochondrial dynamics during oogenesis in *Drosophila*.

### Atg1, Drp1 and Marf collaborate to regulate oogenesis in *Drosophila*

GSCs undergo differentiation and proceed through multiple stages of development termed as egg chambers. Egg chambers undergo further development and they are supplemented with factors necessary during embryogenesis. Previous reports suggest that both Drp1 and Marf are crucial for oogenesis, and their absence leads to detrimental effects on the development of the ovary[6]. Atg1 has been demonstrated to be dispensable for oogenesis as *Atg1KD*, or null mutants, do not disrupt egg development[13]. Our data also suggests that overall egg development is not disrupted in *Atg1KD*. However, the average area of the ovary is significantly reduced in *Atg1KD* (**Figure 4A, B, G and Supplementary Figure 4A, F-I**). Further, the number of GCs (Vasa positive) and the percentage of ovarioles showing development of pre-vitellogenic eggs and vitellogenic egg stages (stage 8 and after) does not change in *Atg1KD* as compared to the controls (**Figure 4H, I**).

**Figure 4.**
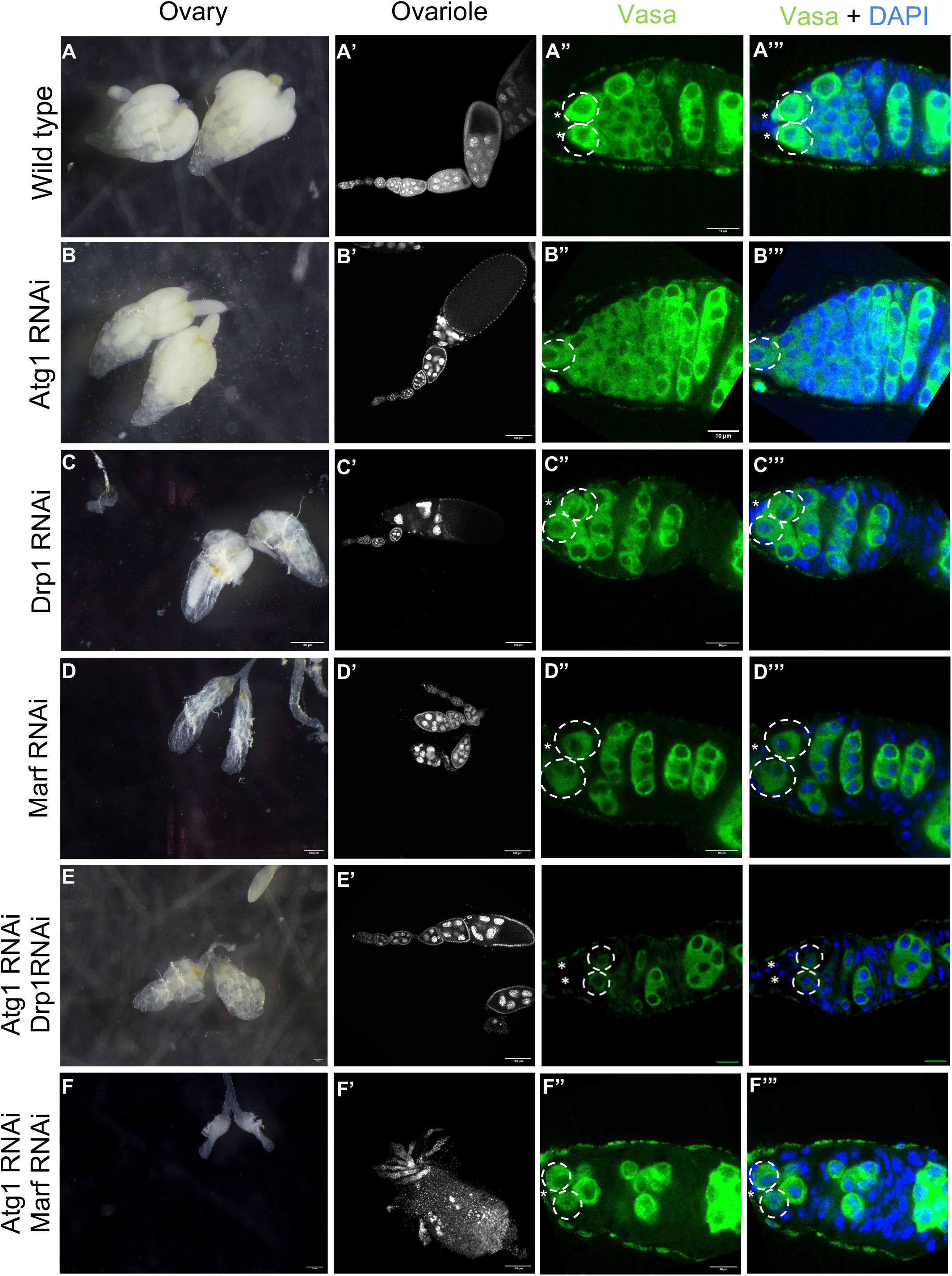
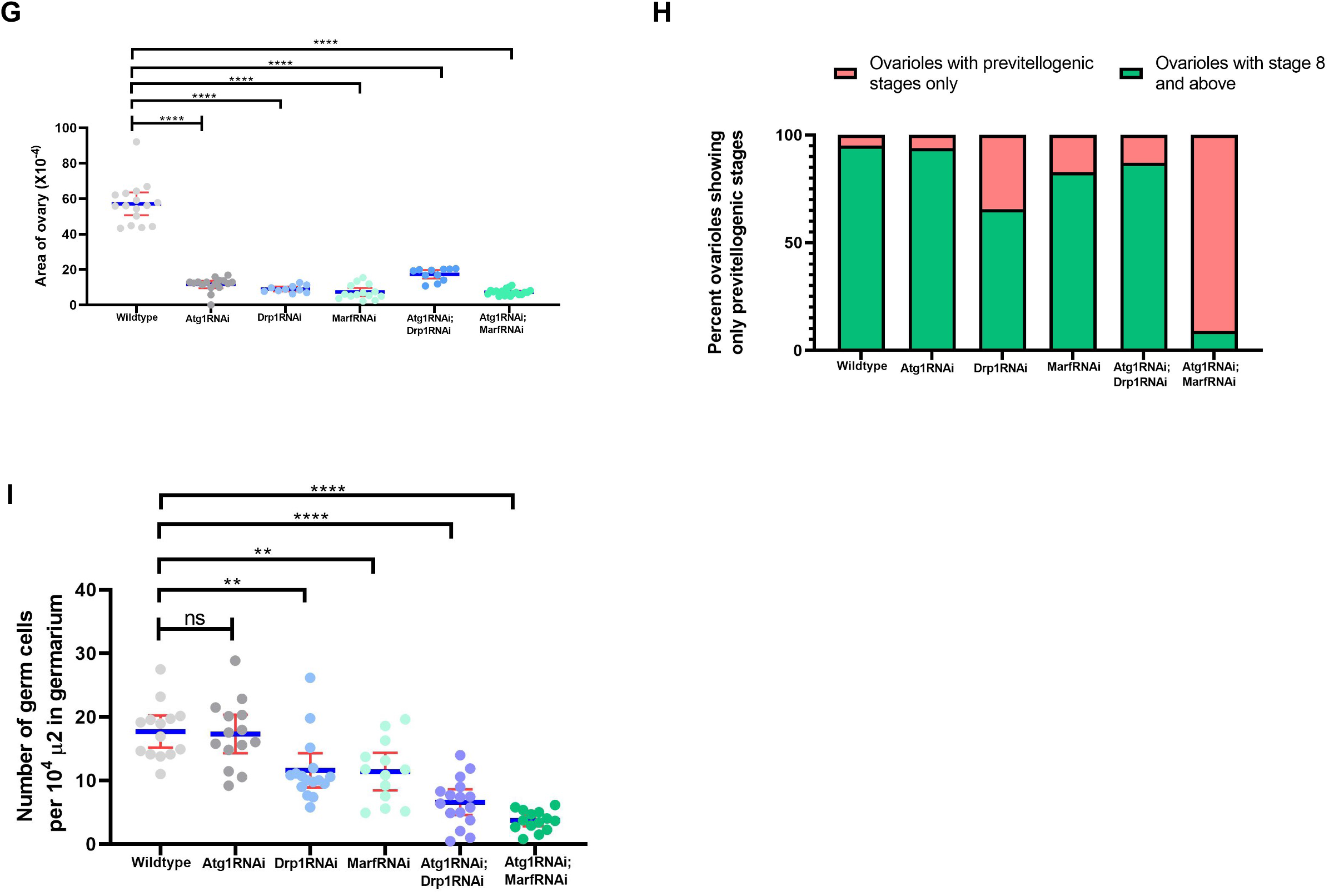
Atg1, Drp1 and Marf collaborate to regulate several aspects of oogenesis. (A-F)Ovaries from wildtype, *Atg1RNAi, Drp1RNAi, Marf RNAi, Atg1RNAi:UASDrp1* and *Atg1RNAi:MarfRNAi* (A’-F’) Ovarioles from wildtype, *Atg1RNAi, Drp1RNAi, Marf RNAi, Atg1RNAi:UASDrp1* and *Atg1RNAi:MarfRNAi*. (A”-F”) germarium immunostained with Vasa (green) to mark the GCs in wildtype, *Atg1RNAi, Drp1RNAi, MarfRNAi, Atg1RNAi:UASDrp1* and *Atg1RNAi:MarfRNAi*. (G) Interleaved scatter plot showing change in overall size of the ovary of wildtype, *Atg1RNAi, Drp1RNAi, MarfRNAi, Atg1RNAi:UASDrp1* and *Atg1RNAi:Marf* (n=10). (H) Stacked bar graph representing the percentage of ovarioles with only pre-vitellogenic and vitellogenic stages in ovary of wildtype, *Atg1RNAi, Drp1RNAi, Marf RNAi, Atg1RNAi:UASDrp1* and *Atg1RNAi:MarfRNAi* (n>100).(I) Interleaved scatter plot showing the change in absolute counts of GCs (Vasa positive cells), quantified in images taken across the central plane of the germaria of ovary of wildtype, *Atg1RNAi, Drp1RNAi, Marf RNAi, Atg1RNAi:UASDrp1* and *Atg1RNAi:MarfRNAi* (n=20). Nuclei is marked in blue (DAPI). Dotted ovals mark the GSCs and asterisk mark the cap cells. Scale bar-10μm. Error bars represent SD in red and the mean is represented in blue. **p < 0.05, **p < 0.01, ****p < 0.0001.

We measured additional parameters of ovarian development such as ovary size, ovariole area, distribution and germ cell differentiation in individual and combined RNAis of *Atg1:Drp1 and Atg1:Marf*. We observed that individual *Atg1, Drp1* and *Marf KD*, as well as combined KD of *Atg1* and *Marf*, led to a significant reduction in the size of the ovary, size of the ovariole and decreased production of egg chambers, both previtellogenic and vitellogenic stages. (**Figure 4B-D’”, F-F’, G-I and Supplementary Figure 4A-I**). Double *Atg1:Drp1 KD* did not exhibit reduction in ovary size and ovariole size (**Figure 4C-C’”, E-E”’**). Strikingly, double knockdown of *Atg1:Marf* show poorly developed ovaries and ovarioles with very few older egg chambers (Stage3 eggs and above) (**Figure 4B-B’, D-D’, F-F’, G, H and Supplementary Figure 4A-C”, E-E” and F-I**). In all controls and *Atg1KD*, the distribution of GSCs (a-spectrin) and GCs (vasa) including 2, 4, 8 and 16-celled cysts was comparable (**Figure 4A”-F’, B-B”’ and Supplementary Figure 4A-E”**). Interestingly, loss of *Drp1* or *Marf* alone exhibited germaria with a significantly less GCs in the germarium (**Figure 4A”-D”**). We observed further reduction in the number of GCs in *Atg1:Drp1* and *Atg1:Marf*. Double KD. *Atg1:Marf* double KD exhibited the strongest GC loss (upto 90%), and this had a detrimental effect on oogenesis. (**Figure 4I, J and Supplmentary Figure 4D-E”’, H and I**). Depletion of GSCs and their differentiated progeny GCs has been previously linked to the failure of oogenesis.

We next asked if GSC maintenance was affected in *Atg1KD* and *Atg1:Marf* double *KD*. We did not consider *Drp1KD* and *Atg1:Drp1* double KD for GSC maintenance assay as Drp1 levels did not alter in *Atg1KD*. Also, previous studies have shown that *Drp1* mutants exhibit GSC loss or gain based on the use of *Drp1* mutants and *Drp1RNAi* and the assay used [6,7]. We expected GSC loss in *Atg1KD* as compared to the controls. Unexpectedly, average GSCs counts were higher in *Atg1KD* at midlife as compared to the controls (**Figure 5A-D’**). Increased fusion of mitochondria was shown to reduce GSC loss due to aging and was observed case of decreased Drp1 or increased Marf activity. *Atg1:Drp1* double KD did exhibit a GC loss phenotype. Interestingly, however, the double KD of *Atg1:Marf* exhibited a dramatic loss of GSCs as early as 1week and a complete loss of GSCs at midlife of the flies. To further assess the GSCs phenotypes in *Atg1KD* and *Atg1:Marf* double *KD*, we measured pMad levels in GSCs. pMad, a self-renewal signal in GSCs is reduced in Marf mutants [7]. We detected a significant loss of pMad intensity in both *Atg1KD* GSCs (**Figure 6A-A’, B-B’ and E**). and *Atg1:Marf* double *KD* suggesting that the reduction in pMad may lead to GSC loss (**Figure 6C-C’, D-D’ and E**). In summary, these data suggest that Drp1, Marf and Atg1 function together to regulate both the maintenance of GSCs and differentiation of GCs during *Drosophila* oogenesis.

**Figure 5.**
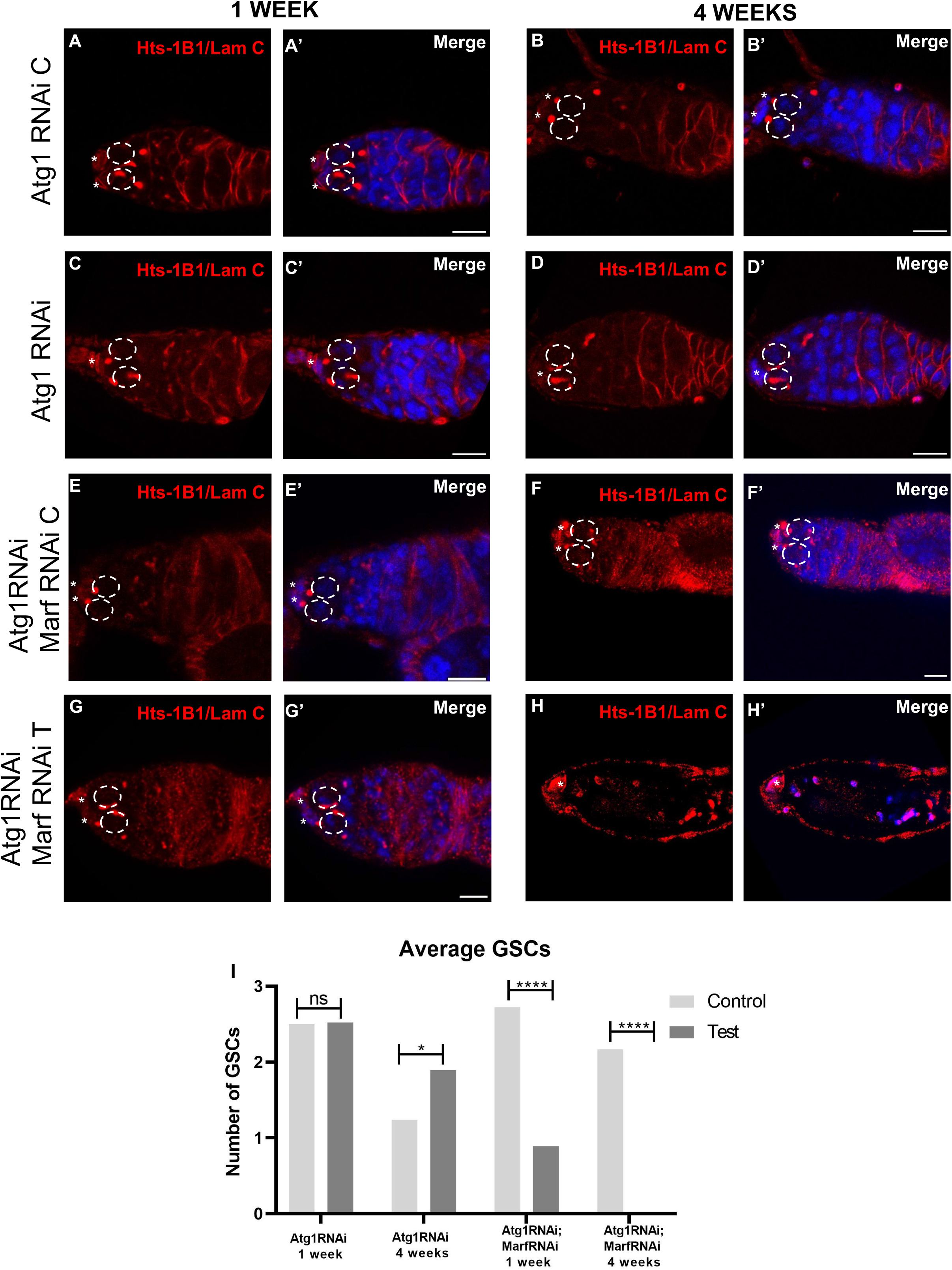
Loss of Atg1 and Marf results in loss of GSCs. (A-H) Representative images for germarium immunostained with hts-1B1 (red) to identify spectrosomes and Lamin C (red) to visualize niche cells in control and *Atg1RNAi* and *Atg1RNAi; MarfRNAi* driven by nosGal4VP16. (I) Stacked bar graph showing the decrease in average GSC number in control and test across 1week and 4 weeks. N=78 for *Atg1RNAi*, N= 36 for *Atg1RNAi;MarfRNAi*, Nuclei is marked in blue (DAPI). Dotted ovals mark the GSCs and asterisk mark the cap cells. Scale bar- 10μm. n=20, **p < 0.05, **p < 0.01, ****p < 0.0001.

**Figure 6.**
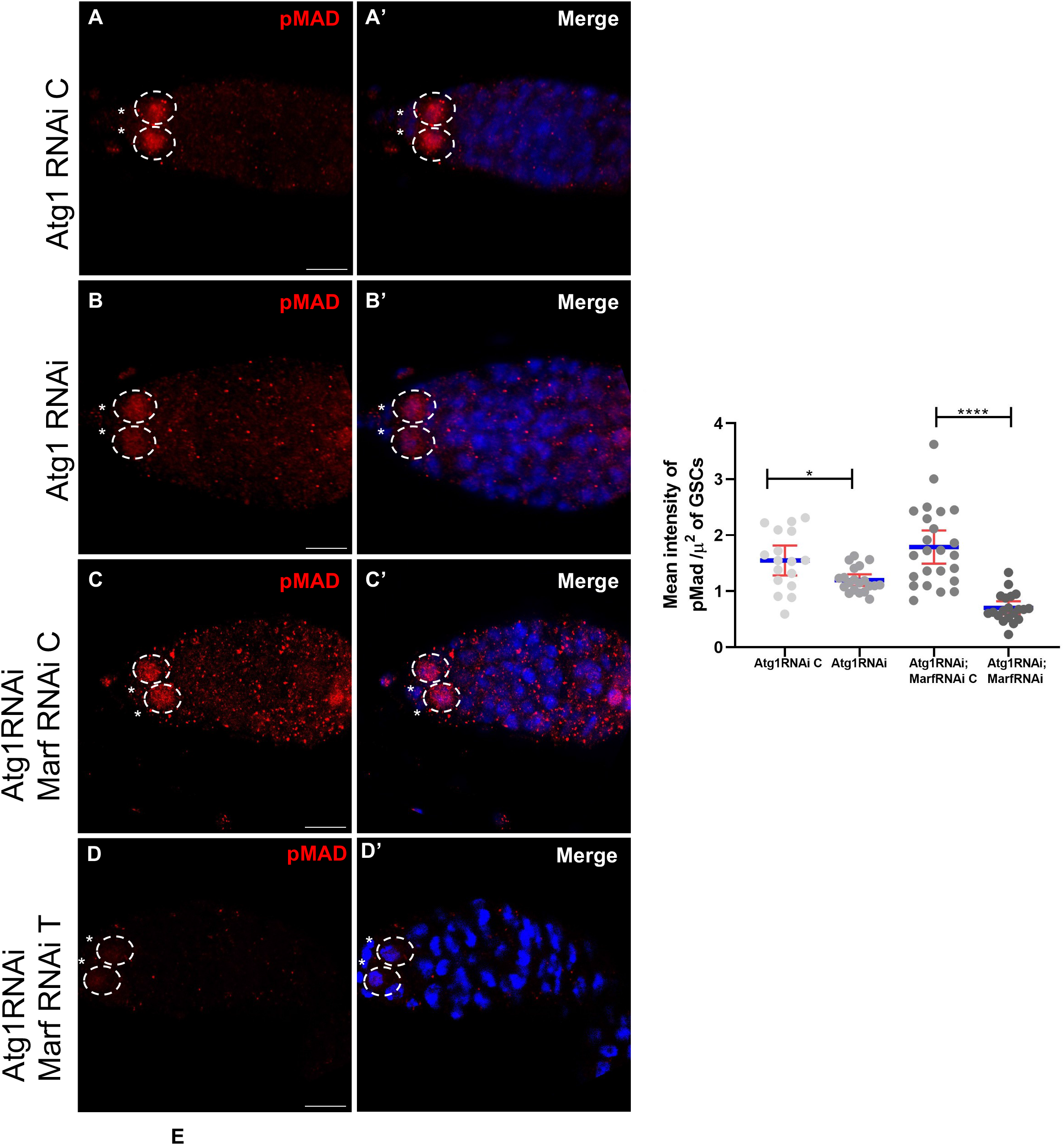
pMad levels reduce in *Atg1RNAi; Marf RNAi*. (A-D) Representative images for germarium immunostained with anti-pMAD in *Atg1RNAi* and *Atg1RNAi; MarfRNAi* driven by nosGal4VP16 and their respective controls. (E) Interleaved scatter graph showing the mean intensity of pMad per unit area in GSCs of control versus test. Error bars represent SD in red and the mean is blue. *n*>25 Nuclei are marked in blue. Dotted ovals mark the GSCs and asterisk mark the cap cells. **p < 0.05, **p < 0.01, ****p < 0.0001

## Discussion

Our work demonstrates that *Autophagy related gene-1 (Atg1*) in addition to its role in the regulation of autophagy (and mitophagy) in *Drosophila*, also regulates mitochondrial dynamics during oogenesis. Atg1 is a core autophagy protein in *Drosophila* that exerts its function through its kinase activity by phosphorylating substrates necessary for the upregulation of autophagy in several tissues. Loss of *Atg1* has been previously reported to disrupt autophagosome formation, and we observed disruption of autophagosome formation in *Atg1RNAi* expressing cells as well as *Atg1-/-* clones in the germarium [3,13]. Additionally, we also observed the accumulation of Ref(2)P in *Atg1KD* germaria, confirming the disruption of autophagy-mediated clearance of protein aggregates. Further, our data also show increased colocalization of Ref(2)P with mitochondria in GSCs and cysts expressing *Atg1RNAi*, indicating disruption of mitophagy. Thus, our data demonstrate that mitophagy in the *Drosophila* ovary is controlled by Atg1, similar to its role in other tissues during different cellular contexts. The mechanism by which Atg1 regulates mitophagy during oogenesis is not understood. Atg1 in yeast and ULK1(the mammalian homologue of Atg1) are known to regulate mitophagy by phosphorylating BNIP3, FUNDC1 and Parkin. However, if Atg1 kinase directly phosphorylates BNIP3, FUNDC1 and Parkin in *Drosophila* during oogenesis needs to be explored.

Strikingly, our data suggest that Atg1 regulates mitochondrial dynamics during oogenesis. Drp1 and Marf expression levels revealed an interesting correlation with the mitochondrial fusion phenotype observed in our study. Drp1 levels remained largely unchanged in GSCs expressing *Atg1RNAi*. It is important to note that in controls, the levels of Drp1 are significantly higher in region 1 as compared to region 2 of the germarium. This difference in Drp1 expression in region 1 Vs region 2 of germanium is disrupted in *Atg1RNAi* (**Supplementary Figure 2A**). Thus, these data suggest that Atg1 influences Drp1 expression. Interestingly though, this did not lead to increased mitochondrial fragmentation. Instead, mitochondria were fused suggesting that fusion phenotype is independent of Drp1 in this context. Marf levels exhibited a significant increase in both GSCs and particularly in the cyst cells, where the mitochondrial fusion phenotype was most prominent. This suggested that the fusion of mitochondria in *Atg1RNAi* is primarily mediated by elevated Marf and not through reduced Drp1. Marf is downregulated in wild-type germline cysts to facilitate mitochondrial selection by enabling the fragmentation of mitochondria. Interestingly, we observed that impaired Drp1 function could cause mitochondrial fusion leading to the formation of “mitochondrial islands” similar to *Atg1RNAi* (**Figure 4**). Thus, it is possible that the mitochondrial fusion phenotype could be due to changes in the ratio of expression Drp1 to Marf rather than changes in their expression. These findings need to be further explored in the context of mitophagy and mitochondrial dynamics during *Drosophila* oogenesis.

The Atg1 mitochondrial fusion phenotype could be rescued by *Drp1^OE,^* suggesting that Drp1 and Atg1 interact genetically. Atg1 has been shown to regulate mitochondrial dynamics in the context of Parkinson’s disease caused by the loss of Pink1/Parkin. Atg1 overexpression (*Atg1^OE^*) rescues the *Pink1/Parkin* mutant-associated fused mitochondrial phenotype by driving mitochondrial fission. Further, it was demonstrated that Atg1OE-mediated fission requires Drp1 as *Drp1* knockdown abrogates the rescue of PD-associated mitochondrial phenotype by *Atg1* overexpression. Interestingly, in another study, the beneficial effects of *Drp1^OE^* prolonging the lifespan of *Drosophila* could be abrogated by *Atg1* knockdown, suggesting that *Drp1* and *Atg1* interact genetically. Drp1 appears to reciprocally regulate *Atg1* mRNA levels. *MarfKD* in *Atg1RNAi* rescues the mitochondrial fusion phenotype of *Atg1* knockdown, indicating that Atg1 also interacts with *Marf* genetically. However, the average mitochondrial area significantly reduced in the double KD of *Atg1* and *Marf*, suggesting that both Atg1 and Marf function together to maintain mitochondrial mass. Both Drp1 and Marf are regulated by phosphorylation, and given that Atg1 is a serine/threonine kinase it is possible that Atg1 phosphorylates Drp1 or Marf or both. However, it is also likely that Drp1 and Marf could be regulated indirectly by Atg1 through other kinases like AMPK, Pink1, and MAPK. These studies need to be conducted to understand the relationship between Atg1, Drp1 and Marf.

Previous studies and our work suggest that Atg1 is not necessary in GCs for oogenesis [13]. This could be due to a redundant function by another kinase, Aduk (Ulk3 homologue in Drosophila) in the absence of *Atg1*. It is also possible that since basal autophagy levels are very low in GSCs loss of *Atg1* is not detrimental [14]. However, it remains to be tested if combined loss of Atg1 and Aduk function is detrimental during oogenesis. We observed a significant reduction in ovary size in *Atg1KD* indicating that Atg1 might control growth of the ovary. A previous study shows that Atg1 is required in follicle cells during oogenesis. Thus, an important question arises “What is the precise role of Atg1 in GCs, including GSCs?”. To address this, we tested if *Atg1* depletion has any effect on GSC maintenance. It was demonstrated that mitochondrial fission promotes GSC loss in the wild-type. Thus, we hypothesized that the depletion of *Atg1* could be beneficial for the maintenance of GSCs and may lead to their longer retention in the GSC niche. Our data support this hypothesis. We observed an increase in average GSC counts at midlife. Although, Atg1 is not necessary for oogenesis per se, it is essential for embryogenesis[13,15]. Embryos with depleted *Atg1* do not complete embryogenesis, and this appears to be partly due to inefficient utilization of lipids [15]. It is not known if the disruption of embryogenesis is due to the inheritance of damaged mitochondria within the developing egg [8].

Drp1 and Marf have been reported to regulate germline stem cell maintenance, differentiation, and ovariole development for the development of vitellogenic eggs and fecundity[6,7]. *Atg1* and *Drp1* double *RNAi* did not enhance the mitochondrial fusion phenotype of Drp1, suggesting that Atg1 acts downstream of Drp1. The double KD of *Drp1:Atg1* exhibited an increase in the average area of the ovary and an increased number of vitellogenic stages as compared to KD of *Drp1* alone, suggesting a rescue of Drp1 phenotype (Figure 4H). However, we observed a significant reduction in the number of GCs double KD of *Drp1:Atg1*, indicating that Drp1 and Atg1 together regulate GSC and GC differentiation. *Marf* and *Atg1* KD led to a dramatic phenotype leading to an arrested growth of ovariole, a drastic reduction in both GSC and GC number. This correlated with a significant reduction in the mitochondrial mass and GSCs were rapidly lost from the niche as early as 7 days. Our data suggest that pMad, one of the self-renewal signals, is significantly decreased in GSCs with *Marf:Atg1* double KD. These data indicate that Atg1 and Marf interact genetically in a complex manner to regulate GSC maintenance through pMad signaling [7]. Thus, our data suggest a complex network formed by Atg1, Drp1 and Marf during oogenesis that connects mitophagy and mitochondrial dynamics in the maintenance of GSCs in *Drosophila*.

## Author Contributions

MA and BVS designed the experiments. MA performed the experiments. BVS and MA wrote the manuscript. BVS edited the manuscript.

## Acknowledgments

We would like to thank Ms. Amruta Nikam for assisting with fly food preparation. Kiran Nilangekar for help with setting up qPCR. We acknowledge the confocal facility at ARI for assistance with imaging. Thanks to members of the Shravage lab for helpful discussions. We would like to thank Developmental Studies Hydridoma Bank, United States, for providing antibodies and plasmid constructs. Thanks to Prof. L. S. Shashidhara and IISER Fly community and Indian *Drosophila* community for providing fly stocks. Dr. Richa Rikhy, IISER Pune, for fly stocks, reagents, and helpful discussions. Dr. Manish Jaiswal, TCIS, Hyderabad for fly stocks, reagents, and valuable comments on the mitochondrial phenotypes. We would also like to thank Dr. P. K. Dhakephalkar, Director, Agharkar Research Institute, Pune, and entire ARI fraternity for support and access to facilities.

## Data Availability Statement

Data available on request from the authors

---

**Supplementary figure 1.**
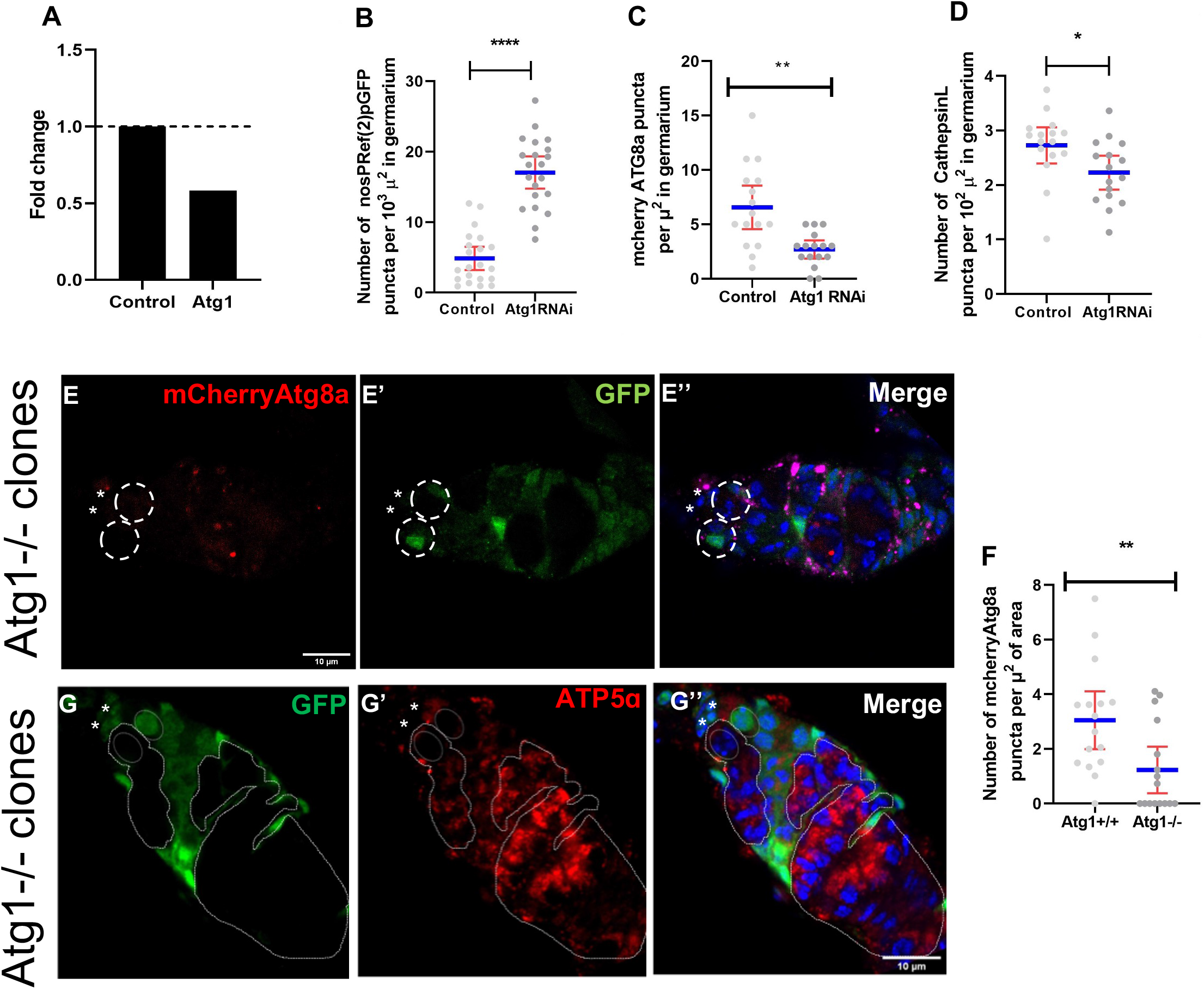
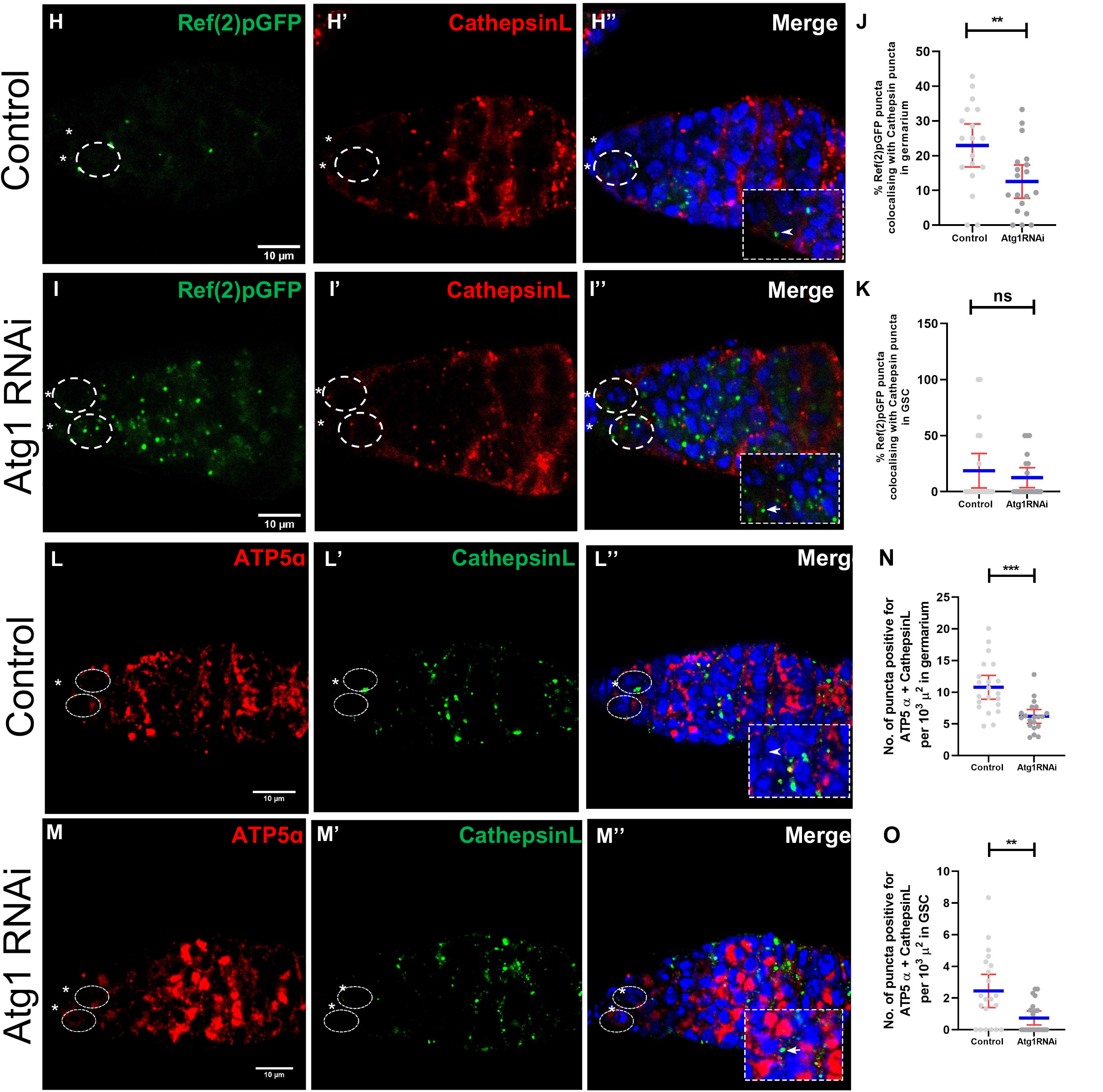
Autophagy is impaired in *Atg1-/-* and *Atg1RNAi*. (A) qPCR analysis shows almost 50% decrease in relative expression of Atg1 in *Atg1RNAi* ovary as compared to control. Interleaved scatter graph representing decrease in autophagosomes (B) Ref(2)PGFP accumulation (C) and Cathepsin L individual counts (D). (E) mCherryAtg8a(red) and CathepsinL (magenta) in *Atg1-/-* germarium outlined and identified by absence of GFP (green). *Atg1-/-* clones are marked with dotted lines. (F) Interleaved scatter graph showing individual puncta counts for mCherryAtg8a in *Atg1* mutant clones. (G-G’) ATP5α (red) staining in Atg1-/-mutant cells are identified by absence of GFP (green) and are marked with dotted lines. (H,I) Control (H-H”) and Atg1RNAi (I-I”) germarium showing Ref(2)PGFP (H’,I’ green) and CathepsinL (H”,I” red) staining. Arrowheads point to colocalized punctae. Inset shows enlarged region of complete colocalization. Arrowheads point to colocalized punctae and arrows indicate Ref(2)PGFP punctae (cargo). (J,K) Interleaved scatter graph representing overall decrease in percentage colocalization between puncta/structures positive for CathepsinL and Ref(2)PGFP in control and Atg1RNAigermarium (G) and GSCs (H). Control (L) and Atg1KD (M) flies immunostained with ATP5α (red) and Cathepsin (green) antibody. Inset shows enlarged region of complete colocalization. (N,O) Interleaved catter graph showing unchanged/ colocalization of Cathepsin-L and ATP5α puncta in germarium(C) and GSCs (D) of Atg1IR and control flies. Nuclei is marked in blue (DAPI). Dotted ovals mark the GSCs and asterisk mark the cap cells. Scale bar- 10μm. Error bars represent SD in red and the mean is represented in blue. n=20, **p < 0.05, **p < 0.01, ****p < 0.0001.

**Supplementary figure 3.**
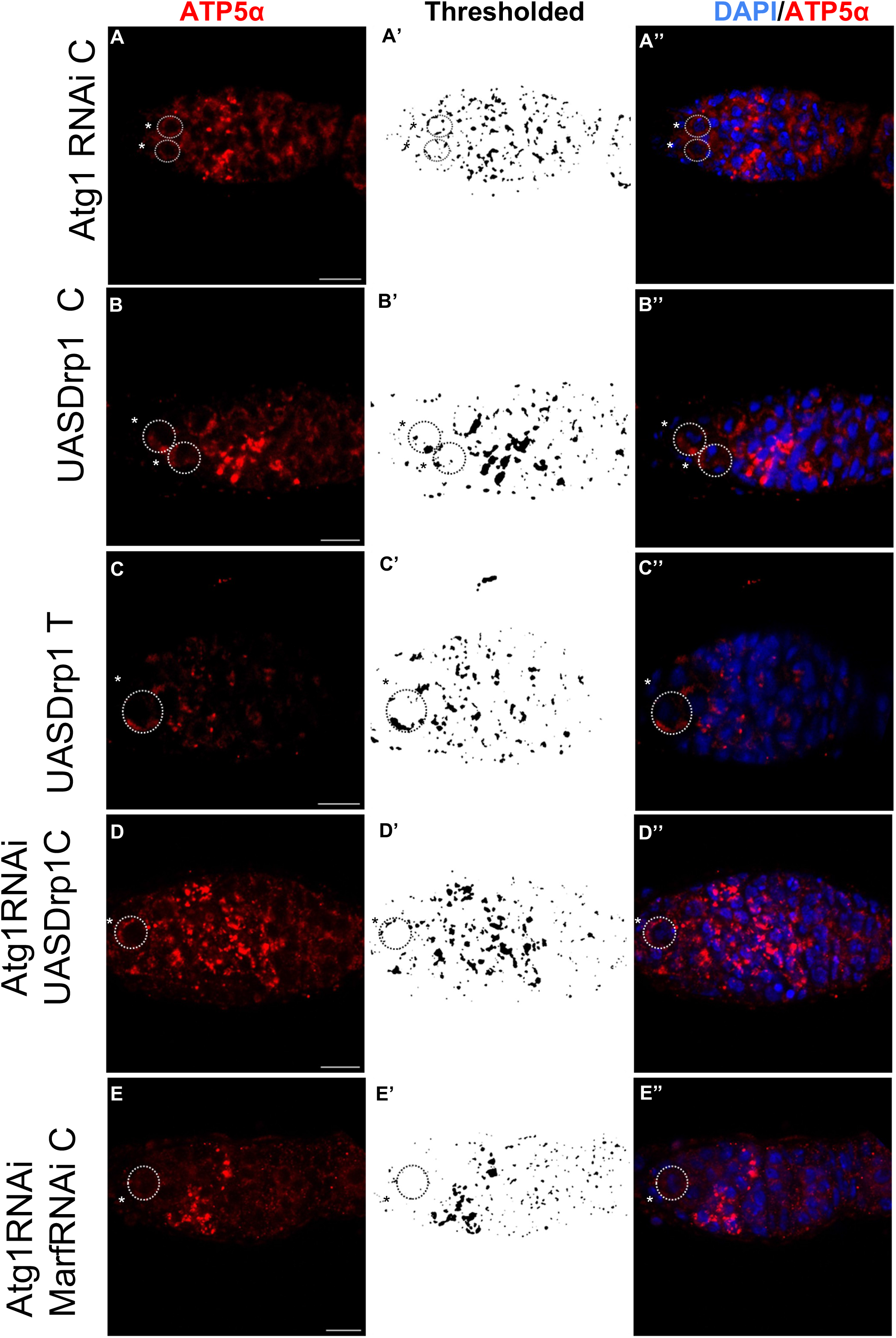
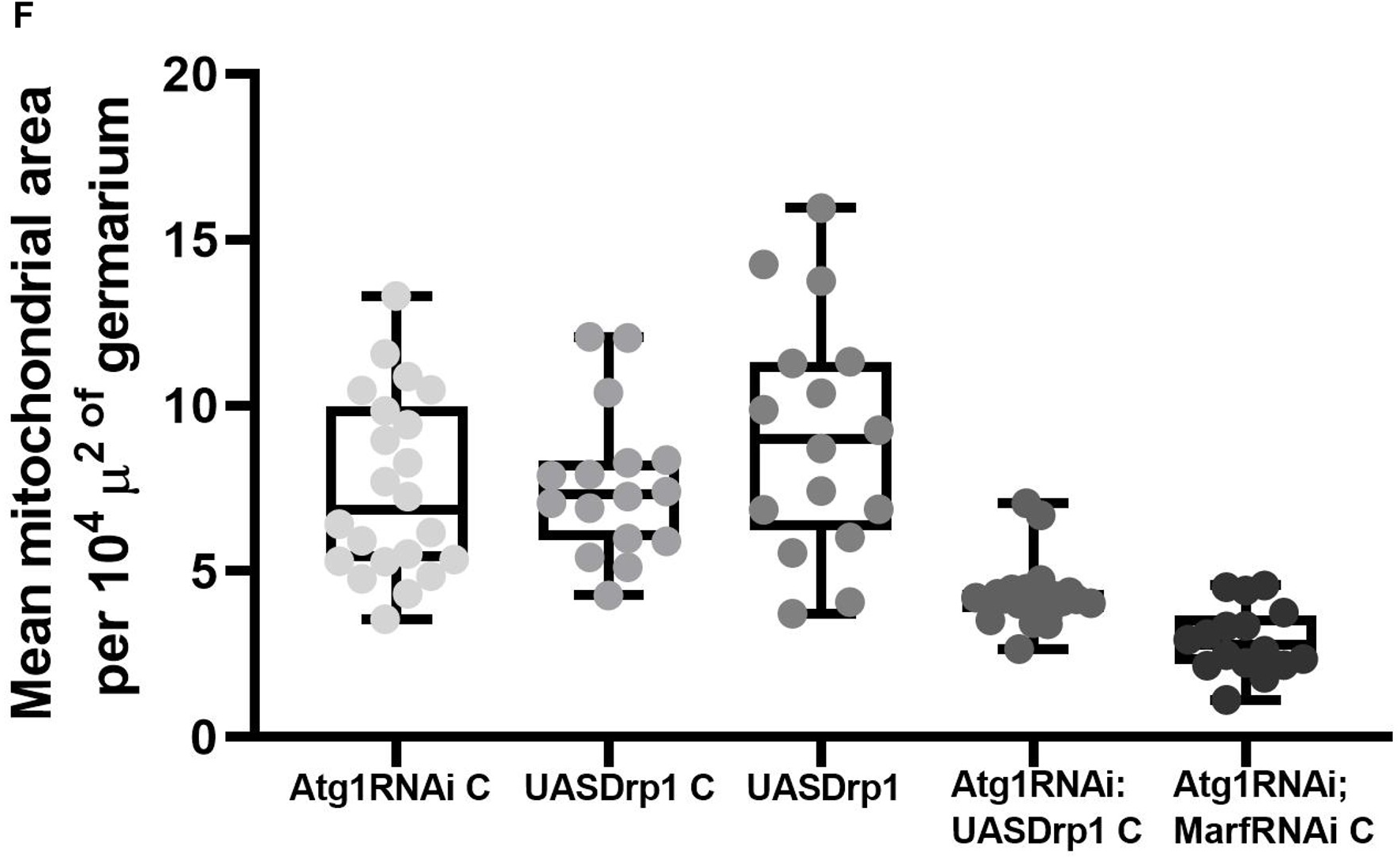
Drp1 and Marf are required for maintaining mitochondrial dynamics. (A) Interleaved scatter plot showing distribution of Drp1 across region 1 and region 2 of control and *Atg1RNAi* germarium. (B) Interleaved scatter plot showing intensity of Drp1-HA in control and *Drp1RNAi* germarium (C) qPCR analysis showing decrease in the relative expression of *Drp1* in *Drp1RNAi* ovaries compared to control. (E) qPCR analysis for relative gene expression for *Atg1* control versus *Drp1RNAi*. Dotted line at 1 represents fold change (F,G) control germaria immunostained for ATP5 Drp1RNAi control (F) and Marf RNAi control (G), thresholded mitochondria in Drp1RNAi control (F’) and Marf RNAi control (G’)and merge with DAPI (F”-G”).Nuclei are marked in blue. Dotted ovals mark the GSCs. Scale bar- 10μm. Error bars represent SD in red and the mean is represented in blue. n=20, **p < 0.05, **p < 0.01, ****p < 0.0001.

**Supplementary figure 5. Mitochondrial homeostasis is crucial for oogenesis**

(A-E”’) Representative brightfield images of whole ovaries dissected from the controls for indicated RNAi. Letter C represents control and T represents test. (*Atg1RNAi C, Drp1RNAi C, Marf RNAi C, Atg1:Drp1RNAi, Atg1:MarfRNAi C*). (A’-E’): Representative images of ovarioles from the controls for indicated RNAi(A”-E”). (A”’-E”’) Germarium of controls for indicated RNAi immunostained for Vasa (green) to mark the GCs. (F) Interleaved scatter plot showing the size of the ovary in controls for indicated RNAi. (n=10) (G) Stacked bar graph representing the percentage of ovarioles with only pre-vitellogenic and vitellogenic (stage 8) stages in controls for the indicated RNAi (n>100). (H) Interleaved scatter plot showing the change in ovariole sizes in wildtype versus RNAi driven by *nosGal4VP16* and their respective controls(n=20). Nuclei are marked in blue. Dotted ovals mark the GSCs. Scale bar- 10μm. Error bars represent SD in red and the mean is represented in blue. n=20, **p < 0.05, **p < 0.01, ****p < 0.0001

**Supplementary figure 2.**
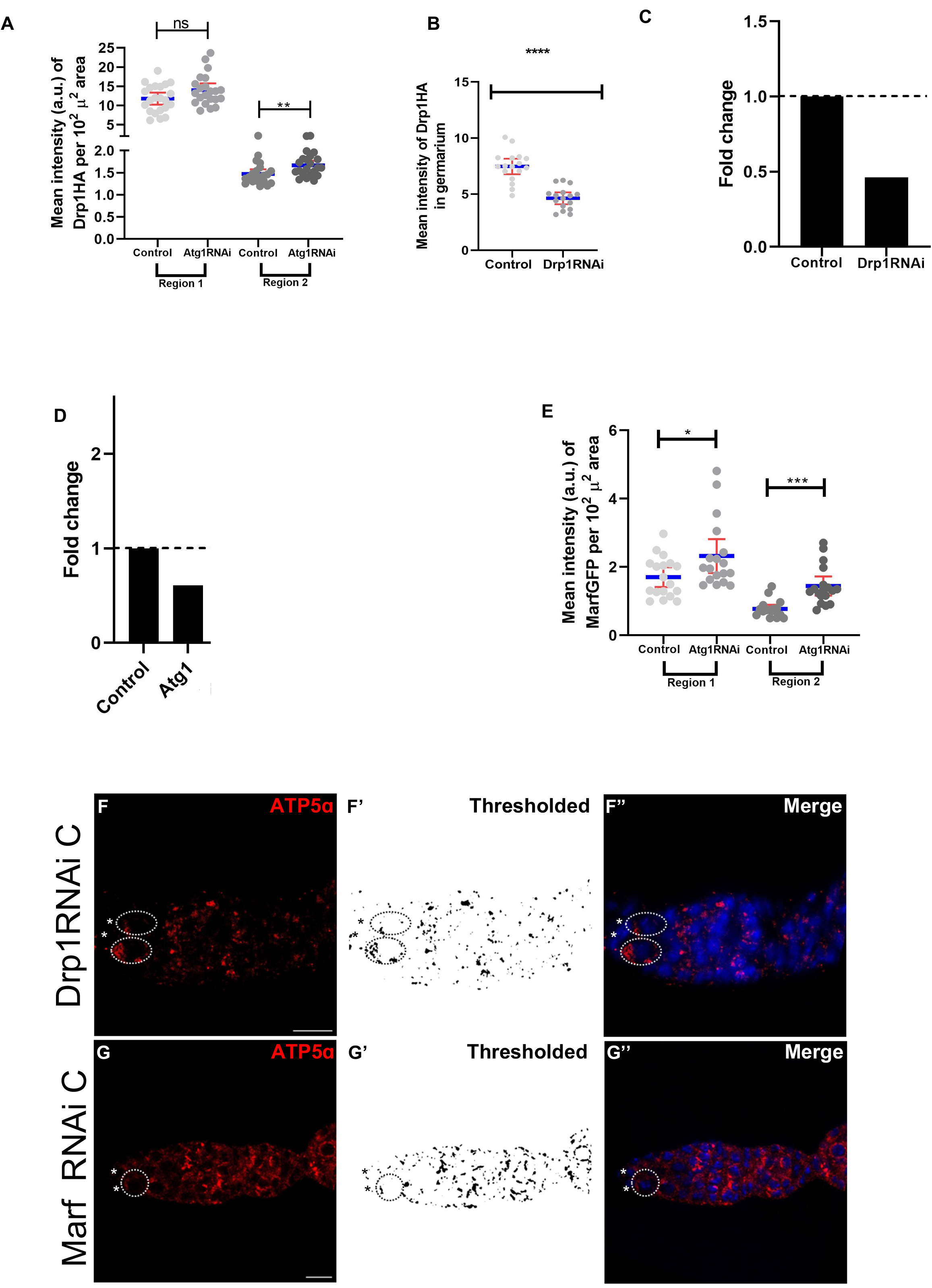
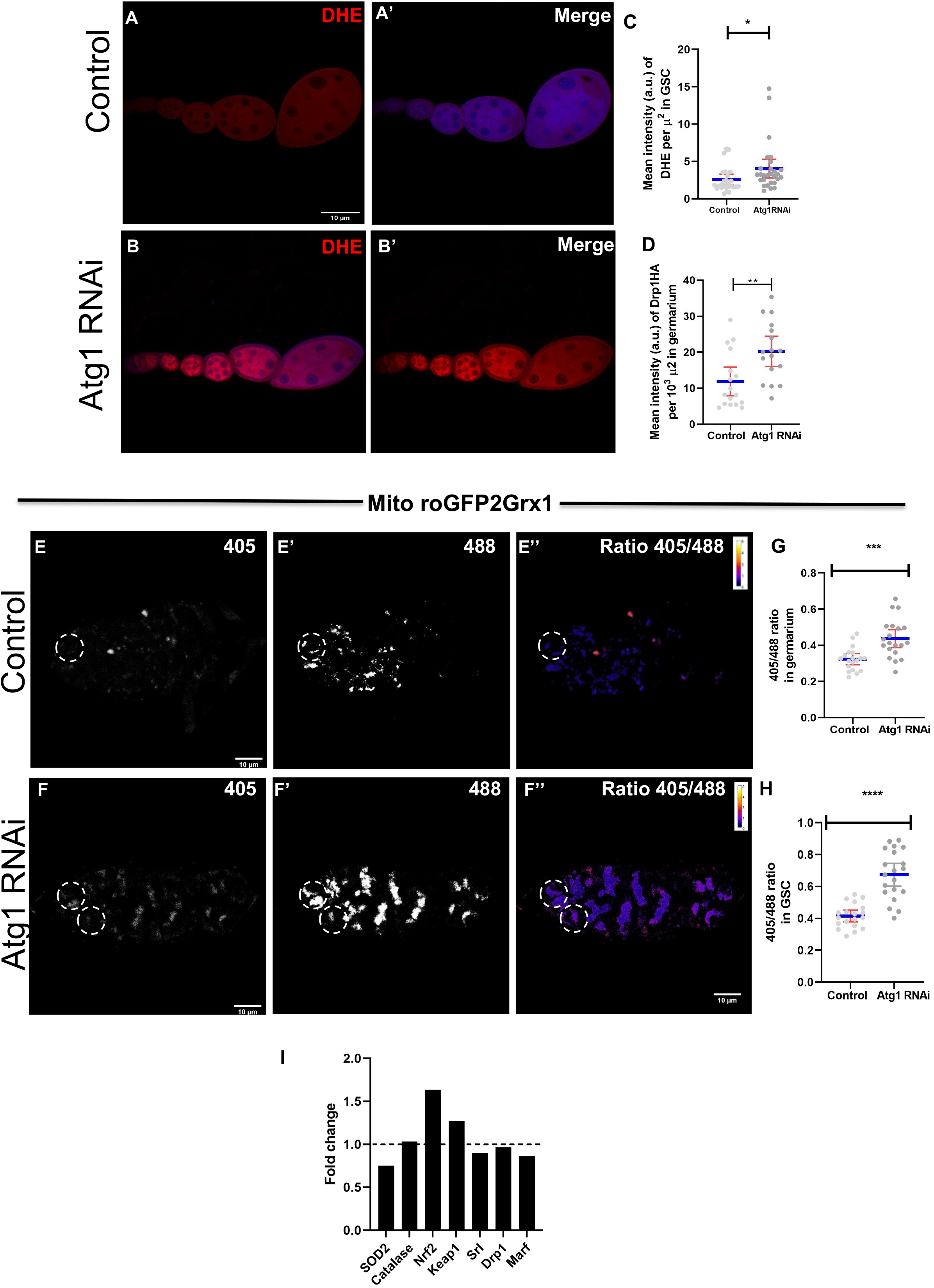
*Atg1RNAi* results in increased ROS. (A,B) DHE (red) staining in control (A) and Atg1KD (B) ovariole. (C,D) Interleaved scatter graph showing mean intensity of DHE in GSCs © and germarium (D) per unit area. (E,F) mito-roGFP2-Grx1 monitors the redox state in the GSC and the germarium. Ratio of emission at 405 and 488 nm obtained as a response of the reporter in control (E) and Atg1KD (F). (G,H) Interleaved scatter graph showing ratiometric shift of excitation in GSC (G) and germarium (H), DR stands for dynamic range. (I) Simple bar graph depicting relative expression levels of various genes viz. SOD2, Catalase, Nrf2, Keap1, Srl, Drp1 and Marf responsible for mitochondrial homeostasis in *Atg1RNAi* compared to control ovaries. Nuclei is marked in blue. Dotted ovals mark the GSCs. Scale bar- 10μm. Error bars represent SD in red and the mean is represented in blue. n=20, **p < 0.05, **p < 0.01, ****p < 0.0001.

**Supplementary figure 4.**
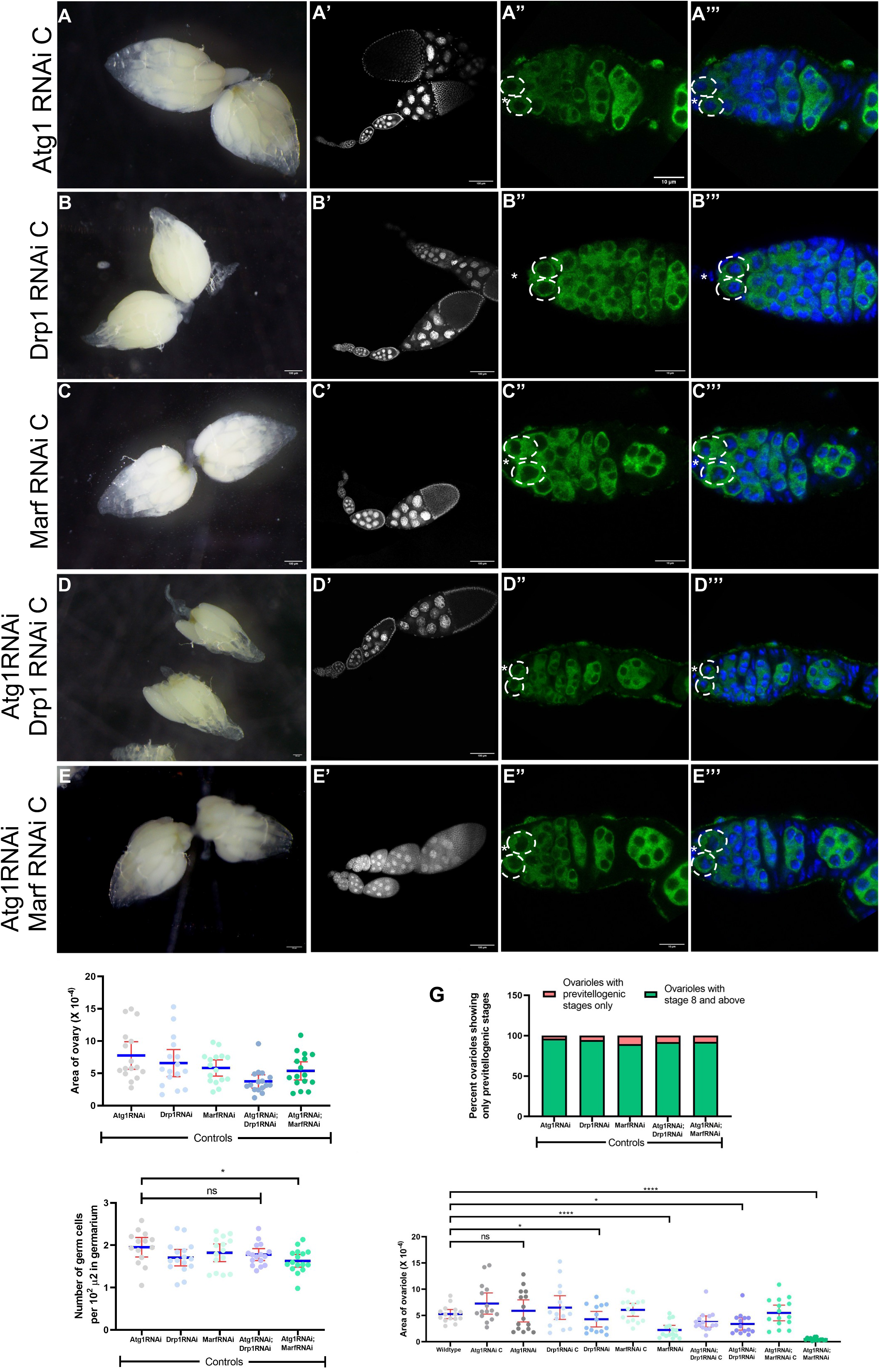
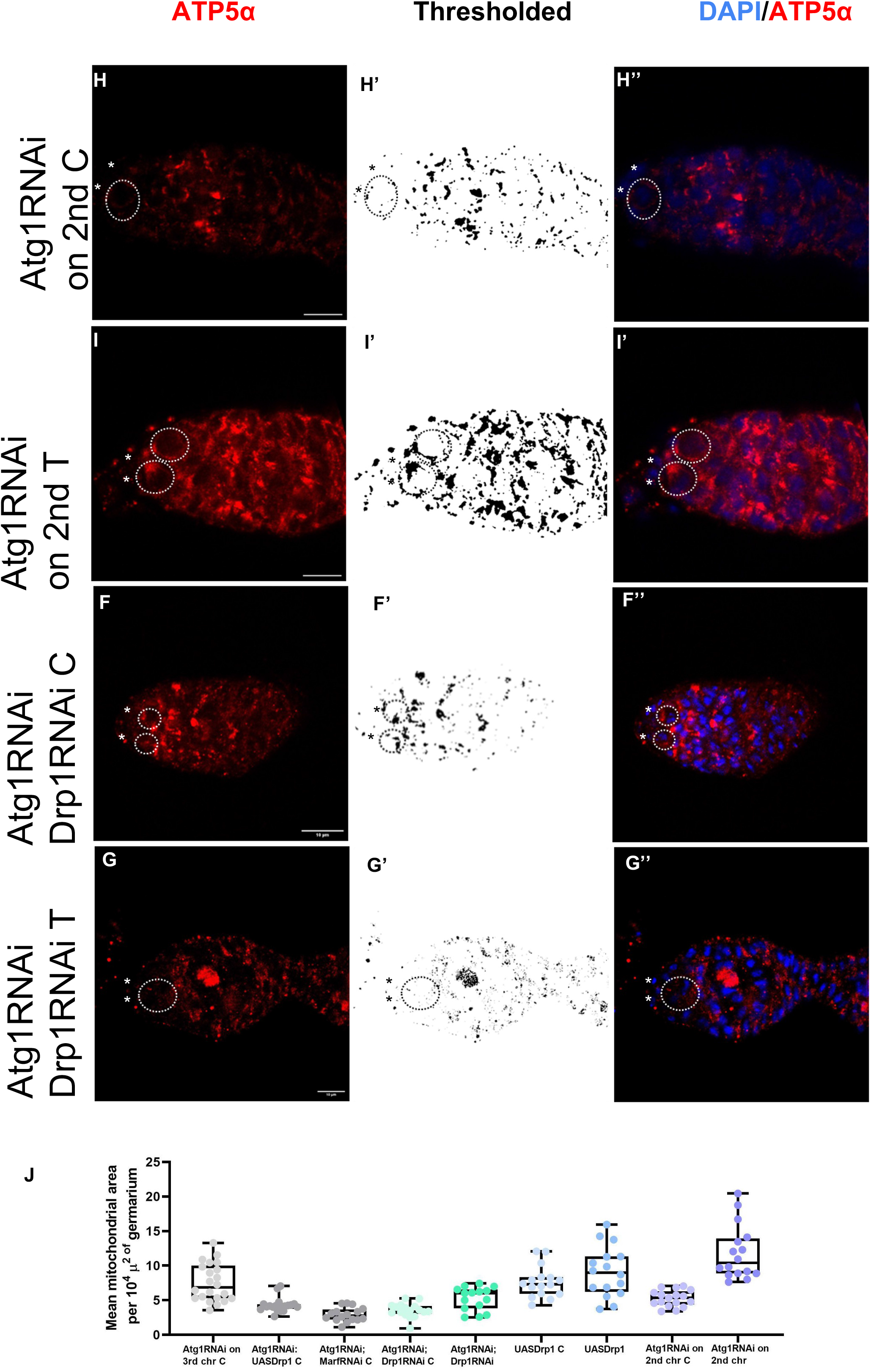
Mitochondrial dynamics is affected by Atg1, Drp1 and MarfKD. (A-I”) Germarium immunostained and quantified for size of mitochondria (red) for the controls and indicated RNAi (*Atg1RNAi C, Atg1:UASDrp1 C, Atg1:MarfRNAi C, Atg1:Drp1RNAi C Atg1:Drp1RNAi T, UASDrp1 C, UAsDrp1 T, Atg1RNAi on 2^nd^ C, Atg1RNAi on 2^nd^ T*). Letter C represents control and T represents test. Representative images of the indicated genotypes for ATP5 (A-I), thresholded mitochondria (A’-I’) and merge with DAPI (A”-I”). (J) Box and whisker plot representing mean area of mitochondria measured per unit area of the germarium. Genotypes are listed on the X axis. Nuclei are marked in blue. Dotted ovals mark the GSCs. Scale bar- 10μm. Error bars represent SD in red and the mean is represented in blue. n=20, **p < 0.05, **p < 0.01, ****p < 0.0001.

## References

[1] M.T. Fuller, A.C. Spradling, Male and female Drosophila germline stem cells: Two versions of immortality, Science (1979). (2007). https://doi.org/10.1126/science.1140861.

[2] K. Nilangekar, N. Murmu, G. Sahu, B. v. Shravage, Generation and characterization of germline-specific autophagy and mitochondrial reactive oxygen species reporters in Drosophila, Front Cell Dev Biol. 7 (2019). https://doi.org/10.3389/fcell.2019.00047.

[3] R.C. Scott, O. Schuldiner, T.P. Neufeld, Role and regulation of starvation-induced autophagy in the Drosophila fat body, Dev Cell. 7 (2004) 167–178. https://doi.org/10.1016/j.devcel.2004.07.009.

[4] R.J. Youle, D.P. Narendra, Mechanisms of mitophagy, Nat Rev Mol Cell Biol. (2011). https://doi.org/10.1038/nrm3028.

[5] G.W. Dorn, C.F. Clark, W.H. Eschenbacher, M.Y. Kang, J.T. Engelhard, S.J. Warner, S.J. Matkovich, C.C. Jowdy, MARF and Opa1 control mitochondrial and cardiac function in Drosophila, Circ Res. (2011). https://doi.org/10.1161/CIRCRESAHA.110.236745.

[6] M. Garcez, J. Branco-Santos, P.C. Gracio, C.C.F. Homem, Mitochondrial Dynamics in the Drosophila Ovary Regulates Germ Stem Cell Number, Cell Fate, and Female Fertility, Front Cell Dev Biol. 8 (2021). https://doi.org/10.3389/fcell.2020.596819.

[7] O. Amartuvshin, C.H. Lin, S.C. Hsu, S.H. Kao, A. Chen, W.C. Tang, H.L. Chou, D.L. Chang, Y.Y. Hsu, B.S. Hsiao, E. Rastegari, K.Y. Lin, Y.T. Wang, C.K. Yao, G.C. Chen, B.C. Chen, H.J. Hsu, Aging shifts mitochondrial dynamics toward fission to promote germline stem cell loss, Aging Cell. 19 (2020). https://doi.org/10.1111/acel.13191.

[8] T. Lieber, S.P. Jeedigunta, J.M. Palozzi, R. Lehmann, T.R. Hurd, Mitochondrial fragmentation drives selective removal of deleterious mtDNA in the germline, Nature. (2019). https://doi.org/10.1038/s41586-019-1213-4.

[9] K.S. Nilangekar, B.V. Shravage, Mitochondrial redox sensor for drosophila female germline stem cells, 2019. https://doi.org/10.1007/7651_2018_167.

[10] K.J. Livak, T.D. Schmittgen, Analysis of relative gene expression data using real-time quantitative PCR and the 2-ΔΔCT method, Methods. 25 (2001). https://doi.org/10.1006/meth.2001.1262.

[11] R.T. Cox, A.C. Spradling, A Balbiani body and the fusome mediate mitochondrial inheritance during Drosophila oogenesis, Development. 130 (2003). https://doi.org/10.1242/dev.00365.

[12] H. Sandoval, C.K. Yao, K. Chen, M. Jaiswal, T. Donti, Y.Q. Lin, V. Bayat, B. Xiong, K. Zhang, G. David, W.L. Charng, S. Yamamoto, L. Duraine, B.H. Graham, H.J. Bellen, Mitochondrial fusion but not fission regulates larval growth and synaptic development through steroid hormone production, Elife. (2014). https://doi.org/10.7554/eLife.03558.

[13] J.M.I.I. Barth, J. Szabad, E. Hafen, K. Köhler, Autophagy in Drosophila ovaries is induced by starvation and is required for oogenesis., Cell Death Differ. 18 (2011) 915–924. https://doi.org/10.1038/cdd.2010.157.

[14] S. Zhao, T.M. Fortier, E.H. Baehrecke, Autophagy Promotes Tumor-like Stem Cell Niche Occupancy., Curr Biol. 28 (2018) 3056–3064.e3. https://doi.org/10.1016/j.cub.2018.07.075.

[15] H. Kuhn, R. Sopko, M. Coughlin, N. Perrimon, T. Mitchison, The Atg1-Tor pathway regulates yolk catabolism in Drosophila embryos, Development (Cambridge). 142 (2015). https://doi.org/10.1242/dev.125419.

